# Bronchoconstriction damages airway epithelia by excess crowding-induced extrusion

**DOI:** 10.1101/2023.08.04.551943

**Authors:** Dustin C. Bagley, Tobias Russell, Elena Ortiz-Zapater, Kristina Fox, Polly F. Redd, Merry Joseph, Cassandra Deering-Rice, Christopher Reilly, Maddy Parsons, Jody Rosenblatt

## Abstract

Asthma is deemed an inflammatory disease, yet the defining diagnostic symptom is mechanical bronchoconstriction. We previously discovered a conserved process that drives homeostatic epithelial cell death in response to mechanical cell crowding called cell extrusion(*1, 2*). Here, we show that the pathological crowding of a bronchoconstrictive attack causes so much epithelial cell extrusion that it damages the airways, resulting in inflammation and mucus secretion. While relaxing airways with the rescue treatment albuterol did not impact these responses, inhibiting live cell extrusion signaling during bronchoconstriction prevented all these symptoms. Our findings propose a new etiology for asthma, dependent on the mechanical crowding of a bronchoconstrictive attack. Our studies suggest that blocking epithelial extrusion, instead of ensuing downstream inflammation, could prevent the feed-forward asthma inflammatory cycle.

## INTRODUCTION

Over 300 million people globally suffer from asthma, with ∼1000 dying from it daily. All asthma attacks are characterized by bronchoconstriction, which causes breathing difficulty, wheezing, and increased airway mucus production. Following severe attacks, asthma patients frequently experience an inflammatory period and airway infection hypersensitivity for weeks to months that can induce further attacks, triggering an inflammatory cycle(*3*). While diverse stimuli can trigger asthma attacks that cause different types of immune responses, the universal life-threatening symptom shared by asthmatics is mechanical bronchoconstriction. In fact, a bronchoconstrictive response to methacholine challenge serves as a standard diagnostic test for asthma(*4*). While current therapies are beneficial, they do not prevent asthma attacks and associated hypersensitivity. Thus, identifying other etiologies driving asthma exacerbations and associated adverse events is a priority.

Epithelia that line the airways provide a protective barrier for the lungs, acting as a first line of defense in innate immunity(*5-8*). Central to providing a tight barrier to the outside world is maintaining constant epithelial cell densities while they turnover at high rates by cell division and death. We discovered a conserved process essential for preserving the barrier as well as epithelial cell number homeostasis called cell extrusion(*1, 2*). Extrusion mechanically links the number of cells dying with those dividing by triggering some cells to seamlessly squeeze out of the layer when they become too crowded(*1, 2, 9*). Given that mild physiological crowding causes homeostatic cell extrusion responses, we postulated that pathological crowding from a bronchoconstrictive attack might trigger so much cell extrusion that it would disrupt the airway epithelial barrier. Destruction of the airway barrier could then lead to the inflammatory period and elevated susceptibility to viral and bacterial infection that typically follow an asthma attack and perpetuate the inflammatory cycle(*10*). Here, we investigate if the mechanics of a bronchoconstrictive event (“asthma attack”) can cause inflammation and damage that could lead to more attacks.

## RESULTS

To test if experimental bronchoconstriction can directly induce airway epithelial cell extrusion, we treated un-primed or immune-primed *ex vivo* mouse lung slices with methacholine (MCH), which triggers airway smooth muscle (ASM) encircling the epithelial barrier to contract(*11, 12*). To prime airways, we used several separate published OVA and HDM immune-priming methods(*13-18*) (Supplemental Fig. 1A, B) known to produce characteristic asthma Th2 inflammatory responses, with increased inflammatory cytokines IL-4 and IL-13 (SFig1C, E) and high mucus production (Supplemental Fig1D, F, and Fig 4A, B). MCH addition caused pronounced bronchoconstriction of mouse lungs immune-primed with either ovalbumin (OVA) or house dust mite (HDM) (as outlined in Supplementary Fig. 1(*13-18*)) but did not significantly affect unprimed airways (Fig. 1 A&B, Supplemental Movies 1-3). Within 15’ of MCH-treatment, the luminal area of small and medium bronchioles reduced dramatically, causing severe airway epithelial crowding (Fig 1B). Importantly, 15’ of bronchoconstriction caused excess airway epithelial extrusion in primed *ex vivo* mouse slices, sparing unprimed controls (Fig 1A, C). To quantify the number of extrusions in response to MCH, we immunostained lung slices with E-cadherin for epithelia, phalloidin for ASM, and DAPI for DNA, scoring for extrusions as cells pinching off apically into the lumen (black data points and red arrowheads) and complete epithelial denuding (blue data points and blue arrowheads, Fig. 1D, C, E). All immune-priming methods induced pronounced bronchoconstriction and excess extrusion in response to MCH, compared to unprimed control mice, with HDM-priming showing the strongest effects (Fig. 1 A, B and Supplemental Movie 2, 3). To test if the amount of constriction correlated with extrusion rates, we used increasing MCH doses on OVA-primed mice and found a tight correlation between the amount of constriction with cell extrusions, with highest doses causing complete denuding of the epithelium (Fig. 1D, E).

**Figure 1.**
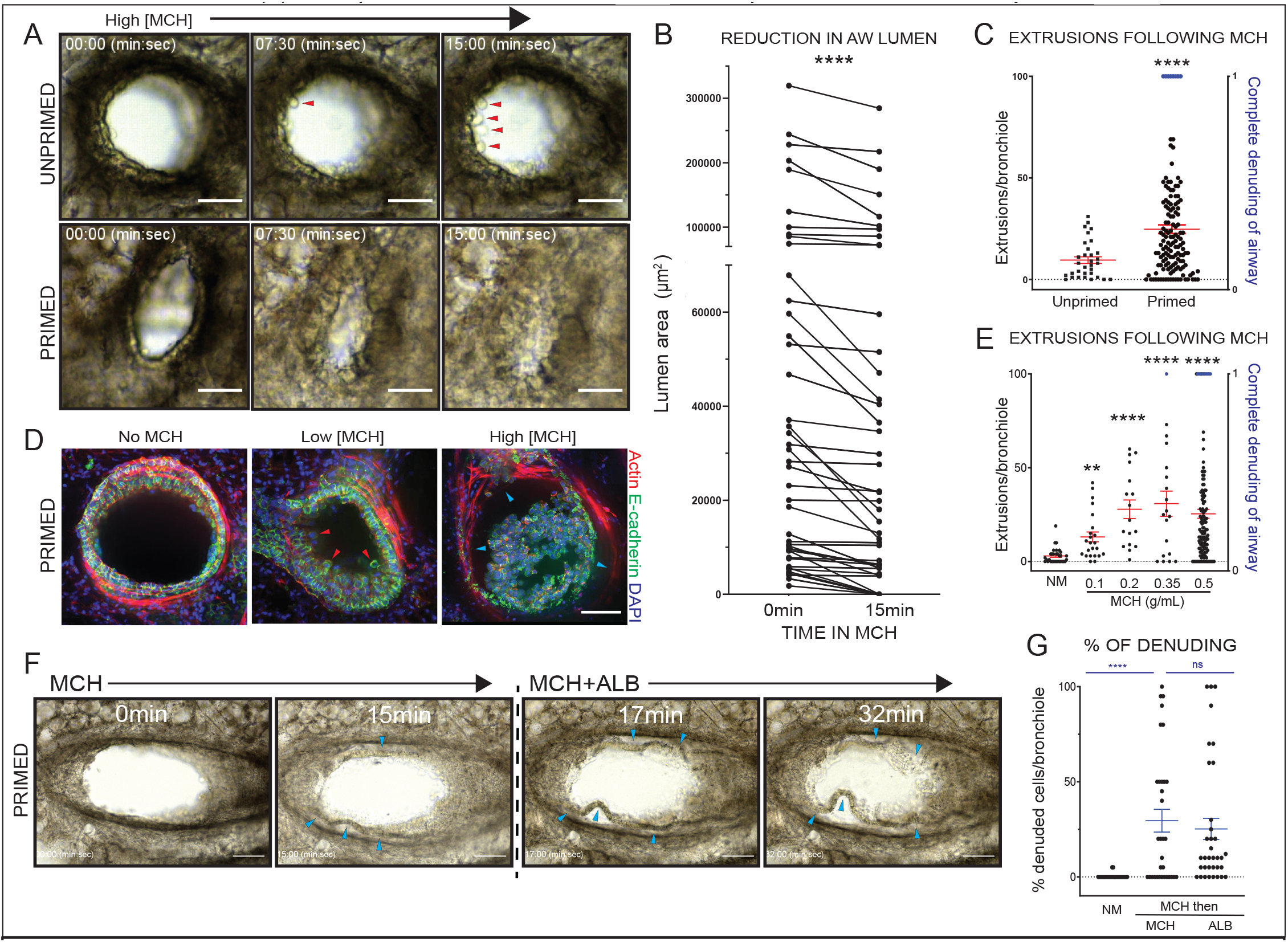
Bronchoconstriction in *ex vivo* lung slices causes excess cell extrusion. (A) Movie stills of 500mg/ml MCH response in unprimed and two week HDM-primed bronchioles (minutes:seconds), with red arrowheads pointing to single-cell extrusions (scale bar=50μm). (B) Constrictions scales with size, measuring the bronchiolar lumen areas before and after MCH treatment in large, medium, and small bronchioles from >5 airways from 5 HDM-primed mice (P****<0.0001 from a Wilcoxon pairs-matched signed rank test). (C) Quantification of extrusions from immunostained *ex vivo* lung slices from unprimed and OVA-primed mice treated with 500mg/ml MCH (P***<0.0005 from an unpaired Mann-Whitney analysis compared to control from >5 lung slices from >10 mice (scale bar=50μm)) in (D), where blue data points represent complete epithelia denuding. (D) Sample confocal projections of bronchioles following 200mg/ml (low), or 500mg/ml (high) MCH, where red arrowheads depict single-cell extrusions and blue ones, epithelial denuding (<100 extrusions on graphs, blue data points). (E) Increasing MCH concentration increases extrusion, (P***<0.0005 and P****<0.0001) from >15 slices per treatment from >5 mice. (F) Movie stills from a 5-week HDM-primed *ex vivo* lung slice treated with 500mg/ml MCH for 15’, and then relaxed with 3.5mM ALB+500mg/ml MCH for 15’, showing epithelial still detach (blue arrows, scale bar=100μm), quantified in (G) with no statistical significance between MCH and ALB from a Mann-Whitney test.

Alleviating airway constriction with the current therapy albuterol, a short-acting ß2-adrenergic receptor agonist, could potentially reverse the excess epithelial extrusion and denuding. However, we found that while albuterol relaxes airways bronchoconstricted for 15 minutes, it did not prevent airway epithelial extrusion and destruction (Fig. 1F&G & Supplemental Movie 4). In fact, albuterol tended to promote more epithelial detachment from the smooth muscle as ASM sprung open, while epithelia remained buckled (Supplemental Movie 4 and Fig. 1F&G, arrowheads). For this reason, we quantified the percent epithelial cell denuding per bronchiole, rather than extrusions (Fig. 1G). Thus, albuterol does not prevent destruction of the airway lining following a bronchoconstrictive attack. Moreover, increased denuding we noted could potentially impede airway repair.

We next investigated if canonical extrusion inhibitors could prevent bronchoconstrictive airway damage. We previously discovered that crowding-induced live cell extrusion requires the stretch-activated channel Piezo1 to trigger production of the bioactive lipid sphingosine 1-phosphate (S1P) that binds the S1P_2_ receptor to activate Rho-mediated actomyosin contraction needed to seamlessly eject a cell from the monolayer(*1*),(*19*) (schematic, Fig. 2A). We find that extruding airway epithelial cells specifically upregulate S1P, the limiting signal for extrusion, suggesting that a conserved extrusion pathway operates in mouse airways (Fig. 2 B). While Piezo1, the stretch-activated channel (SAC) we identified as critical for activating extrusion in response to crowding in MDCKs and zebrafish(*1*), is expressed in mouse airways, it is important to note that they also express the transient receptor protein (TRP) channels TRPA1, TRPV1, and TRPM8 frequently upregulated in unmanageable asthma(*20*) that could act similarly to trigger extrusion (Supplemental Fig. 2). Notably, immune priming caused Piezo1, TRPV1, and TRPM8 to become more vesicular and widespread (Supplemental Fig. 2). Thus, to inhibit Piezo1 as well as other potential TRP channels, we used gadolinium hexahydrate chloride (Gd^3+^) a generic SACs and TRP inhibitor. Additionally, we blocked S1P production with sphingosine kinase 1 (SKI II) and 2 (K-145) inhibitors and the receptor S1P_2_ with the antagonist JTE-013 during MCH-induced bronchoconstriction and quantified extrusion and denuding. All inhibitors dramatically decreased extrusions following MCH (Fig. 2C, D), however, only Gd^3+^ and SK inhibitors blocked epithelial sheet denuding as well (Fig. 2C, E). It is not clear why only Gd^3+^ and SK inhibitors block detachment, however, it suggests that blocking extrusion upstream in the pathway offers greater protection of the airway epithelium.

**Figure 2.**
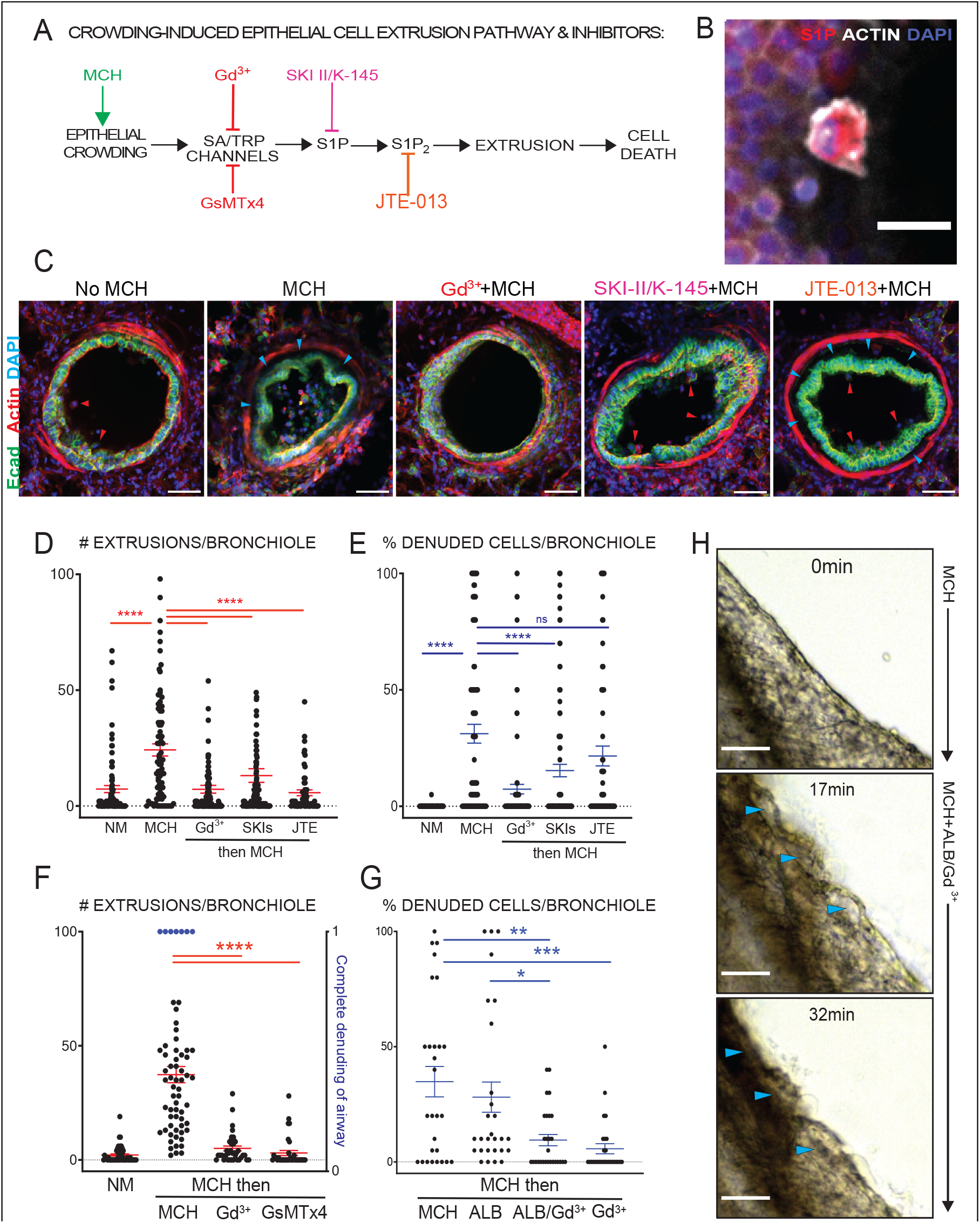
SAC and S1P inhibitors block extrusion caused by bronchoconstriction. (A) Canonical crowding-induced epithelial cell extrusion pathway, indicating where each inhibitor acts within the pathway. (B) Confocal projection of an extruding cell immunostained for sphingosine 1-phosphate (S1P), phalloidin for f-actin, and DAPI (scale bar=25μm). (C) Confocal projections of *ex vivo* lung slices from 5-week HDM-primed mice with no MCH (n=9), MCH (n=9), or pre-treated with Gd^3+^ (n=9), sphingosine kinase inhibitors SKI II&K-145 (n=9), or S1P_2_ antagonist JTE-013 (n=4), before adding MCH (scale bar=50μm), quantified in (D), as extruded cells (red arrows) and (E) as % epithelial denuding (blue arrows) per bronchiole (P****>0.0001 from a Kruskal-Wallis test, from >4 slices/mouse from >4 mice). Quantification of OVA-primed lung slices treated with Gd^3+^ or GsMTx4 *after* 15min of MCH challenge, where black dots represent extrusions per bronchiole and blue dots represent complete epithelial denuding from >5 mice (F) or percent denuding (G) (P*<0.05, P**,0.001, P***<0.0005 and p****<0.0001 from a Mann-Whitney test, from >4 slices/mouse from >4 mice). (H) Movie stills from an HDM-primed mouse lung slice pretreated with MCH for 15’, following ALB (3.5mM)+MCH (0.5g/ml), with arrowheads pointing to areas of epithelial denuding that reattach by 32’ (scale bar=50μm). Note that albuterol alone does not prevent epithelial destruction.

We next tested if we could inhibit extrusion in a more clinically relevant way—after the onset of bronchoconstriction and in the presence of albuterol. Thus, we triggered bronchoconstriction with MCH for 15 minutes and then added SAC inhibitors in the presence of MCH. Remarkably, Gd^3+^ administered after bronchoconstriction dramatically reduced extrusion and epithelial denuding (Fig. 2F-H). Additionally, the peptide inhibitor, GsMTx4 that blocks Piezo1, TRPC1, and TRPC6(*21*), also blocks extrusion when added 15 minutes after bronchoconstriction (Fig. 2F). Importantly, albuterol did not impair the ability of Gd^3+^ to block epithelial extrusion/denuding following 15 minutes of MCH-induced bronchoconstriction, nor did Gd^3+^ affect bronchodilation by albuterol (Fig. 2G,H, Supplemental Movie 5, 6). Our videos show that Gd^3+^added to albuterol allows the airway epithelia to reattach to the underlying smooth muscle as it relaxes, thereby preventing denuding (Fig 2H, blue arrowheads, and Supplemental Movie 5, 6). Again, it is unclear why blocking upstream in the extrusion pathway allows epithelial reattachment but suggests that it prevents secretion of matrix proteases predicted to be required during the extrusion process.

The ability for Gd^3+^ to prevent bronchoconstriction-induced airway epithelial extrusion and destruction suggested that it may prevent the inflammation that typically follows an asthma attack. To test this hypothesis, we challenged live HDM-primed mice with albuterol and inhibitors of extrusion, compared to control and MCH alone, and assayed for extrusions, denuding (30min post MCH), and inflammation (24hrs post MCH) by H&E staining. We identified 50mg/ml MCH (1/10 the concentration used in *ex vivo* experiments) as the lowest dose that would trigger bronchoconstriction and extrusion without causing undue harm and stress to mice, delivering it in increasing amounts, as administered with patients, to ensure that no mice were hyperresponsive (Fig. 3A, schematic). We then administered Gd^3+^ or sphingosine kinase inhibitors (SKIs) during MCH challenge to test if blocking early defined steps in the extrusion pathway we identified as protective in our ex vivo assay would prevent subsequent inflammation. SKIs can be repeatedly administered safely to mice intranasally(*22*) and we find that intranasal instillation of 10 μM GdCl_3_ can be given to mice for five consecutive days (Supplemental Fig. 3B) or once a week for 3 weeks (Supplemental Fig. 3C) without causing masses, inflammation, or noticeably affecting mouse health and behavior (Supplemental Movies 7&8). Because albuterol does not impede the ability of Gd^3+^ to block extrusion (Fig. 2 E-H and Supplemental Movie 5&6) and is necessary for opening airways in patients, we used albuterol with Gd^3+^ treatment in all our live mice studies to reduce total numbers of mice. Histology slices 30 minutes after MCH challenge with or without albuterol resulted in high numbers of extrusion (red arrows) and denuded bronchioles (blue outline, Fig. 3B-D). Yet, addition of Gd^3+^ with albuterol preserved airway epithelia, similar to those with no MCH challenge (NM) (Fig. 3B-D). The damage incurred during bronchoconstriction with or without albuterol resulted in pronounced immune cell infiltration by 24 hours, compared to immune-primed unchallenged (NM) bronchioles (Fig. 3E&F). Strikingly, inhibiting extrusion with gadolinium or SKIs (with albuterol) reduced the inflammatory response to levels seen in control immune-primed, unchallenged (NM) bronchioles (Fig. 3E&F). Similar results were obtained with OVA-primed mice, even when Gd^3+^ is delivered after the methacholine ramp-up (Supplemental Fig.4). Thus, preventing bronchoconstriction-induced extrusion, particularly with gadolinium, prevents the inflammation that typically follows an attack.

**Figure 3.**
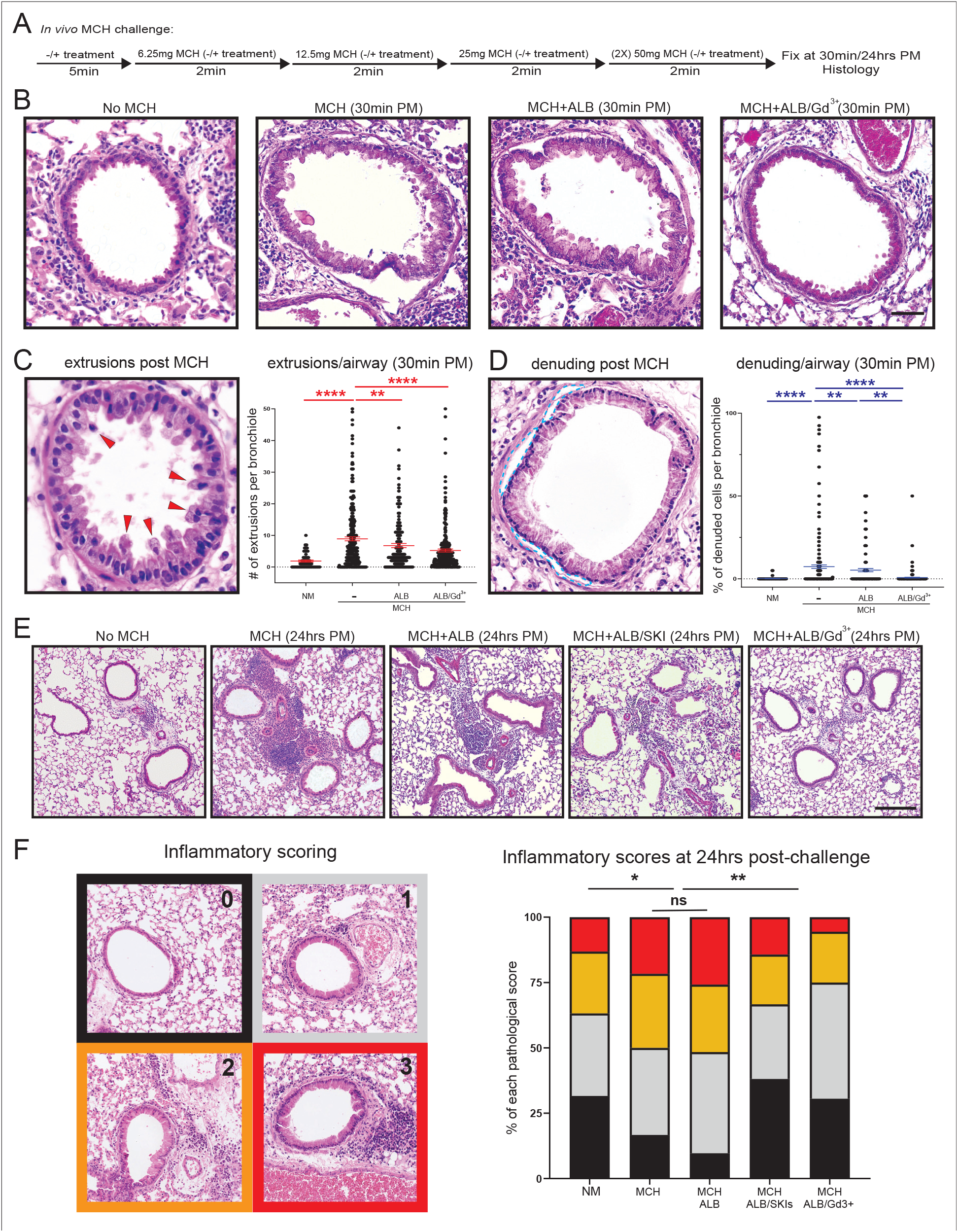
SKIs and gadolinium block extrusion and inflammation following a bronchoconstrictive attack in live mice. (A) Schematic of live MCH challenges ± 5min pre-treatment with extrusion inhibitors before increasing MCH ± inhibitors. (B) H&E sections from no MCH (NM) (n=4), MCH, or treated with MCH (n=5), MCH+ALB (n=5), or MCH+ALB/Gd^3+^ (n=6) at 30min post MCH, with the number of extrusions and percent denuding per bronchiole quantified in (C&D), with representative images depicting single-cell extrusions (red arrows) and bronchiole denuding (blue outline) (P**,0.001, P***<0.0005 and p****<0.0001 from a Mann-Whitney test). (E&F) H&E sections 24hrs post MCH-challenge to observe immune response from no MCH treatment (n=4), or MCH alone (n=5), MCH+ALB (n=5), MCH+ALB/SKI (n=3), or MCH+ALB/Gd^3+^ (n=5) with representative images shown in (E) and quantification of airway immune cell infiltration using pathological scores, defined by colors and numbers in (F) (P*<0.05 and P**<0.001 from a Chi-squared test).

While asthma typically damages the distal airways, most asthma patients experience significant difficulties with excess mucus secretion from the primary airways. Immune priming with OVA or HDM predictably increased mucus production and secretion, as measured by qRT-PCR (Supplemental Fig. 1D&F), Muc5AC immunostained confocal projections (Fig. 4A), and Periodic Acid Schiff (PAS) staining of primary airways (Fig. 4C & Supplemental Fig. 1D&F)(*13, 14, 23-26*). Live imaging of primary airways loaded with Wheat Germ Agglutinin-350 (WGA-350) to label mucus from three-week HDM-primed mice revealed that MCH induces rapid mucus secretion with bronchoconstriction, as shown by the loss of blue fluorescence (Fig 4B and Supplemental Movies 9&10). Surprisingly, Gd^3+^ dramatically reduced mucus secretion in primary airways in response to our *ex vivo* MCH challenges (Fig. 4B, Supplemental Movie 11). While our WGA-350 assay is not a very quantitative assay, due to uneven expression of mucus from immune priming and WGA loading, we found approximately 90% of control primary airways secrete mucus, where only ∼26% of GD3+-treated ones do. *In vivo*, we saw a similar response in both OVA and HDM primed mice. PAS staining from HDM-primed mice challenged with MCH for ∼30’ showed bulk mucus secretion with large mucus globules squeezing out apically, which was not prevented by albuterol (Fig. 4C&D), whereas Gd^3+^ treatment with albuterol dramatically reduced mucus secretion (Fig. 4C&D). Similar results were found from OVA-primed mice (Supplemental Fig. 5).

**Figure 4.**
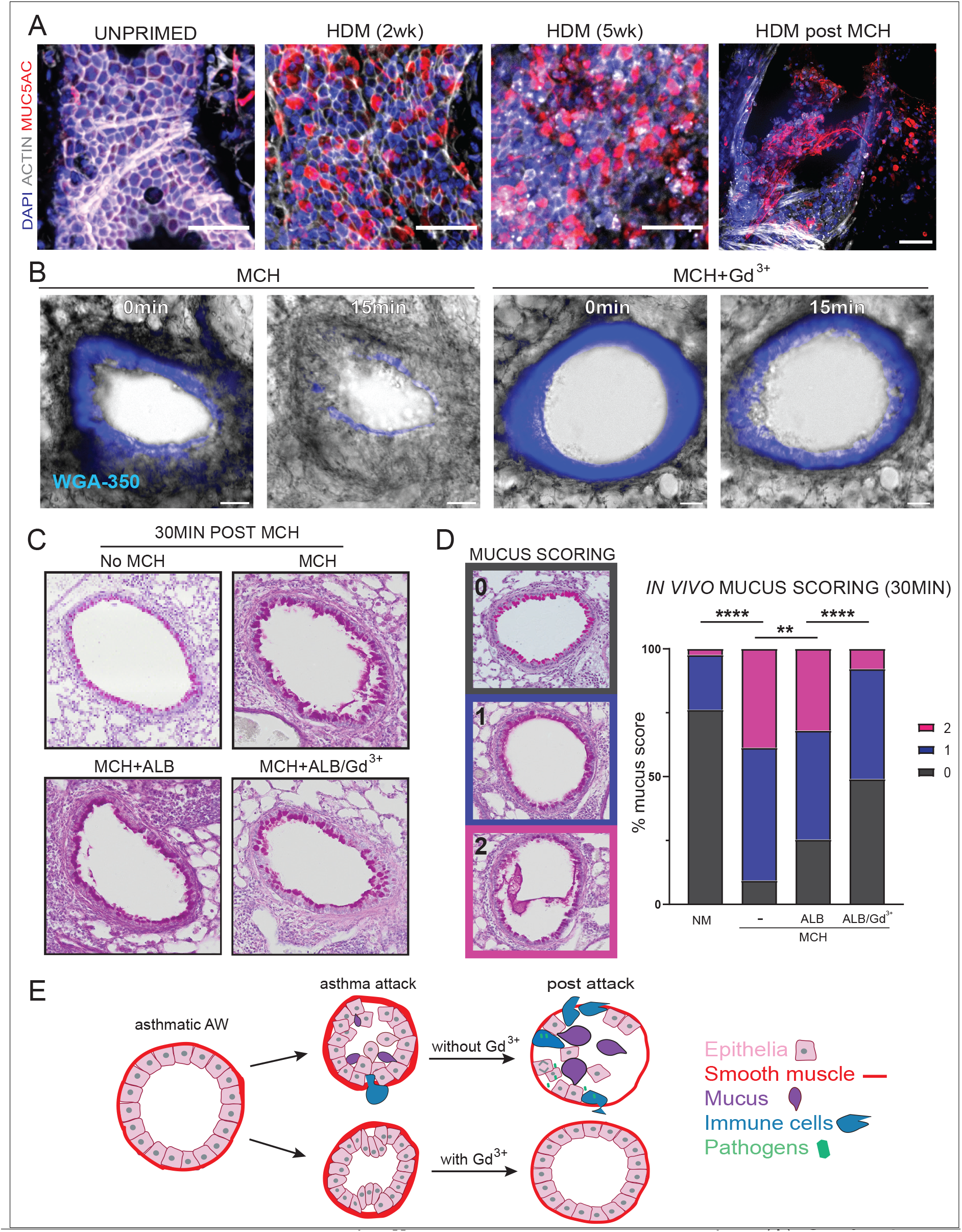
Gadolinium blocks mechanically-induced mucus secretion. (A) Confocal projections of Muc5AC before and after two and five weeks of HDM-priming (scale bar=50μm). (B) Movies stills from 3-week HDM-primed *ex vivo* slices incubated with 10μg/mL WGA-350 to label mucus then either treated with 0.5g/mL MCH alone ± 10μM Gd^3+^Cl for 15mins, noting that WGA is retained in the epithelium, rather than secreted with gadolinium. (C&D) Representative PAS (mucus) staining 30’ following a MCH challenge ±ALB or ALB/Gd^3+^, which was quantified using a color/number scoring in (D) with P**,0.001 and p****<0.0001 from Chi-square tests from at least 5 mice per group. (E) Model describing how the mechanics of asthmatic bronchoconstriction cause excessive crowding-induced mucus secretion and epithelial cell extrusion, which that results in epithelial damage and inflammation. Inhibiting extrusion early in the pathway can prevents all of the pathological consequences of an attack.

## DISCUSSION

Here, we present a new etiology for asthma. Whereas most asthma studies have focused on the inflammatory signaling associated with asthma, our work suggests that the inflammation and mucus secretion result from the mechanics of bronchoconstriction on airway epithelium. We find that bronchoconstriction causes pathological airway epithelial crowding, leading to so much cell extrusion that it destroys the barrier, resulting in the typical post-attack inflammation (Fig. 4E). Because epithelia act as the first line of defense against pathogens and toxins, epithelial barrier disruption could also cause infection hypersensitivity, until airways have repaired. Infections and inflammation could then lead to more bronchoconstrictive attacks, triggering the asthma inflammatory cycle(*27*). Importantly, albuterol treatment, standardly used by asthma patients to relax airways, does not prevent airway epithelia destruction, mucus secretion, or inflammation following an asthma attack. However, blocking the extrusion pathway with gadolinium or S1P inhibitors following bronchoconstriction preserves epithelial integrity, dramatically dampening the inflammatory response. Remarkably, gadolinium also blocks mucus secretion from primary airways, suggesting crowding-activation of calcium channels mechanically induces bulk mucus secretion. Our studies define the extrusion pathway in controlling the downstream symptoms of an asthma attack. Admittedly, our studies do not address if Piezo1 or TRP channels specifically control extrusion in response to bronchoconstriction. Thus, future studies will need to determine which channels respond to different asthma triggers. A variety of TRP channels are not only mutated in many cases of poorly controlled asthma cases but are activated by numerous stresses that trigger asthma attacks, including pollution, smoke, and cold, suggesting that several channels may be relevant (*20, 28-30*). Thus, Gd3+ has the advantage of generically blocking extrusion upstream in the extrusion pathway. Additionally, transient use of Gd3+ used has the added benefit that it may be used practically, once an attack occurs without apparent side-effects in mice (Supplemental Fig. 3) but its safety will need to be tested in human airways. Gadolinium not only serves to establish a role for mechanically-induced extrusion in driving asthma symptoms but also suggests that targeting extrusion upstream in the pathway will be the most effective approach to prevent downstream symptoms. Thus, if gadolinium is not safe in humans, development of other early extrusion inhibitors may provide therapeutic benefit.

Because blocking extrusion upstream in the pathway with gadolinium preserves the airway barrier following a bronchoconstrictive attack, it may also prevent the airway smooth muscle remodeling associated with wound healing(*31*) and S1P production(*22, 32, 33*) that causes future attacks. Thus, preventing the mechanical damage caused by an asthma attack could pave the way to a new generation of therapies that stop the whole asthma inflammatory cycle, rather than treating only the downstream symptoms. Moreover, excess smooth muscle constriction may underlie other inflammatory syndromes, especially those linked with cramping, such as irritable bowel syndrome or inflammatory bowel disease, yielding new approaches to these unsolved medical problems.

## MATERIALS AND METHODS

### Animal models and immune-priming methods

Mice at six-eight weeks of age where immunogen-sensitized in the following ways, by adapting and optimizing several protocols (S Fig.1A). ***OVA-priming:*** For intraperitoneal injections, 8mg/mL albumin from chicken (OVA) (Sigma – A5503) was absorbed into an equal volume of Imject® Alum (Thermo Fisher Scientific – 77161) before injecting 200 μl intraperitoneally (i.p.) injected into Balb/c mice, as stated in protocol in SFig. 1A. Mice were then challenged with aerosolized OVA (8 mg/mL for 15min), as indicated in in SFig. 1A for each protocol. All experiments were done within 48hrs of the last OVA challenge. ***House Dust Mite (HDM):*** 6-8 week old C57BL/6J mice were anesthetized with isofluorane and administered 25 μg (total protein) of HDM (Citeq Biologics, Groningen, The Netherlands) in 25 μl PBS intranasally 5 times per week for 2 or 5 weeks as shown in SFig 1B. Analysis was performed 24 hours after the final challenge. We used mice primed as in (OVA A) for Figs 1C-D&E, 2F&G, as in (OVA B) for Supplemental Figs. 4&5 and Supplemental Movie 3, as in (HDM A) for Figs. 1A&F, 2C-E, 3A Supplemental Movies 2&4-6, and as in (HDM B) for Figs. 1B, 3, 4A-D, Supplemental Fig. 2, and Supplemental Movies 3, 9-11.

All animals were maintained under specific pathogen–free conditions and handled in accordance with the Institutional Committees on Animal Welfare of the UK Home Office Animals (Scientific Procedures) Act 1986. All animal experiments were approved by the Ethical Review Process Committee at King’s College London and carried out under license from the Home Office, UK. All mice were humanely handled, according to an IACUC license 16-08007 from the University of Utah and a Program Project License P68983265 to Jody Rosenblatt and P9672569A to Maddy Parsons at King’s College London.

### Precision-cut ex vivo lung slices (PCLSs)

*Ex vivo* lung slices were obtained from mice, within 48hrs of their last allergen priming challenge, adapted from the protocol of (*34*). Briefly, mice were humanely killed by CO_2_ inhalation followed by cervical dislocation or exsanguination via the femoral artery. The chest cavity was opened and the trachea carefully exposed, where a small incision was made to accommodate the insertion of a 20Gx1.25 in canula (SURFLO I.V. catheter). The lungs were inflated with 2% low melting agarose (Fisher – BP1360) prepared in HBSS+ (Gibco – 14025) before lungs, along with the heart and trachea, were excised, washed in PBS, and the lobes separated. Individual lobes were then embedded in 4% low melting agarose and solidified on ice. 200micron thick slices were cut on a Leica VT1200S vibratome and washed and incubated in DMEM/F-12 medium supplemented with 10% fetal bovine serum (FBS) and antibiotics. The *ex vivo* lung slices remained viable (MCH-reactive) for at least one week after isolation (data not shown).

### Methacholine (MCH) treatment and live imaging

*Ex vivo* lung slices were experimentally treated with increasing dosages of MCH (acetyl-β-methylcholine chloride, Sigma A2251), from 100mg/mL to 500mg/mL in HBSS+ solution. MCH is a non-selective muscarinic receptor agonist clinically used to test the severity of asthma in patients. Lungs were incubated in HBSS+ in 24 well plates at 37°C with MCH for 15-30 minutes, filmed at 15-30sec intervals using a Life Technologies EVOS FL Auto microscope to measure bronchoconstriction in response to MCH. For lumen reduction measurements in response to MCH, the lumen area (the “dead” or empty space in the airway lumen) at time=0min and at time=15min of MCH treatment was measured using Fiji software from live imaging stills. In the wheat germ agglutinin (WGA) experiments, *ex vivo* lung slices were pre-incubated with 10μg/mL of WGA AlexaFluor350 (Invitrogen W11263) per manufacturer’s instructions at 37°C before MCH treatment and live imaging as above. In some experiments, 10μM gadolinium (III) chloride hexahydrate (Sigma G7532) and/or 3.5mM albuterol (Sigma PHR1053), was used either before or 15 minutes into a MCH challenge. The SIP inhibitors at 30μM SKI II (Sigma S5696) and K-145 (Merck 5091060001), or JTE-013 (Sigma J4080) were added at the beginning of a MCH challenge. The *ex vivo* lung slices were fixed in 4% paraformaldehyde overnight at 4°C before immunostaining.

### Immunofluorescence and imaging of fixed PCLS

PFA fixed *ex vivo* lung slices were incubated for one hour at room temperature in blocking solution: PBS containing 0.1% triton X-100, 0.1% sodium azide, and 2% bovine albumin (BSA), before incubating overnight at 4°C at 1:100 in blocking solution for all primary antibodies used: rabbit anti-E-Cadherin (24E10 – Cell Signaling 3195), rabbit anti-Piezo1 (Novus NBP-78537), mouse anti-S1P (Echelon Biosciences ZP300), mouse anti-Muc5AC (Abcam ab3649), rabbit anti-TRPV1 (Alomone ACC-030),rabbit anti-TRPV4 (Alomone ACC-034), rabbit anti-TRPA1 (Alomone ACC-037), rabbit anti-TRPM4 (Alomone ACC-044), and rabbit anti-TRPM8 (Alomone ACC-049. *Ex vivo* lung slices were washed 3 × 30 minutes in PBS+0.5% Triton X-100) before incubating overnight at 4°C overnight with: 1:100 Alexa Fluor 488 goat anti-rabbit (Thermo Scientific - A11008) or anti-mouse (A32723) IgG, Alexa Fluor 568 goat anti-rabbit (A11011) or anti-mouse (A11004) IgG, or Alexa Fluor 647 goat anit-rabbit (A32733) or anti-mouse (A21235) IgG + 1:250 Alexa Fluor 488, 568, or 647 Phalloidin (Thermo Scientific – A12379, A12380, A22287, respectively). Slices were washed 2 × 30 minutes in PBS+0.5% Triton X-100, stained with 1:1000 DAPI in PBS for 20 minutes, mounted in ProLong Gold (Invitrogen P36930), and imaged on a Nikon Eclipse Ti2 spinning disc confocal microscope with a 20X or 40X objective.

### Live mice MCH challenge with and without treatments

Mice at the end of their immune-priming protocol were placed in a 6.5-quart Hefty® bin that was fitted with an Aerogen® Pro Nebuliser System (AG-AP6000-XX) that produces 2.5-4.0μm volume mean diameter aerosolized particles, adapted from (Bevans et al., 2017). Mice were given two-minute challenges of increasing concentrations of MCH (6.25, 12.5, 25, then 2 × 50 mg/mL), resting in fresh air for 2 minutes between challenges, with albuterol (3.5mM) and/or gadolinium (10μM), and/or SKIs (30 μM) given in the last three challenges with the highest MCH concentration of 50mg/mL. Mice were humanely killed at 30’ or 24hrs post MCH challenge (-/+ treatments).

### Histological analysis and quantification

At the end of their treatments, mice were humanely killed by CO_2_ inhalation followed by cervical dislocation or exsanguination via the femoral artery. The lungs were inflated with 10% neutral buffered formalin (NBF), excised, and fixed in NBF overnight at room temperature, followed by another overnight room temperature incubation in 70% ethanol. The large left lung, and the cranial, medial, and caudal lobes of the right lung were excised and embedded in paraffin. Using a keratome, 3×5μm thick serial slices were made per slide, 3 slides per lung, each slice made at 20-50μm deeper intervals, and stained with hematoxylin and eosin (H&E) or periodic acid-Schiff (PAS). Histological grades depicted in Fig.3 and Supplemental Fig. 4 were used to determine immune cell infiltrate (H&E) and mucus phenotypes (PAS) in Fig.4 and Supplemental Fig. 5, with at least 30 airways spanning multiple lung lobes from at least five mice analyzed for each group. H&E stained lung slices were also used to determine if the number of extrusions and percent denuding per bronchiole, with five mice analyzed per group (Fig 3B&C).

### Quantitative Real-time PCR analysis

To confirm the induction of a Th2 inflammatory response from our immune priming protocols. The transcript levels of IL-4, IL-13, and Muc5AC (Thermo Fisher TaqMan probes – cat. # 4331182, 4331182, and 4331182, respectively) were measured by quantitative real-time PCR from cDNA transcribed (Thermo Fisher SuperScript™ cat. # - 18091050) from total RNA isolated from whole lung extracts (Qiagen – RNeasy Plus – cat. # 74034) and mRNA expression analysed by a comparative Ct method.

### Statistical analysis

To compare sample means, we used a non-parametric Mann–Whitney U test (Glantz, 2012). For airway lumen constriction measurements in response to MCH treatment, a Wilcoxon pairs-matched signed rant test was used. To compare more than two groups, we used a Mann–Whitney U test, with the Kruskal-Wallis test or the Holm-Sidak adjustment for pairwise comparisons. For categorical data we used the Chi-Square test to reject the null hypothesis with the Yates correction when analyzing two populations and two categories. For analysis of categorical data, when at least 20% of the groups presented frequencies lower than 5 for a given variable, we used the Fisher Exact test.

## ACKNOWLEDGEMENTS

We thank the Rama Krishnan and Alexandra Locke for help with initial experiments and advice for our paper. We thank Saranne Mitchell, Carlos Pardo Pastor, Michael Redd, and Teresa Zulueta-Coarasa for helpful comments on our manuscript.

## CONTRIBUTIONS

J. R. designed experiments, interpreted, analyzed data, and wrote the manuscript. D.C.B., T. R. and K.F. designed experiments, interpreted data, and performed most live and fixed imaging experiments throughout the paper. P. F. R. and M. J. assisted with mouse priming, imaging, and quantification of imaging results of many of the experiments. C.R. & C.D.R. consulted on experiments and pathology sections. E.O.Z provided HDM-primed mice and helped with imaging of ex vivo slices. All authors edited the manuscript.

## DATA AND MATERIALS AVAILABILITY

All data are available in the main text or the supplementary materials.

## SUPPLEMENTARY MATERIALS

Supplementary Figures S1 to S5; Movies S1 to S11.

