## Supplemental Figures for "Bronchoconstriction damages airway epithelia by excess crowding-induced extrusion"

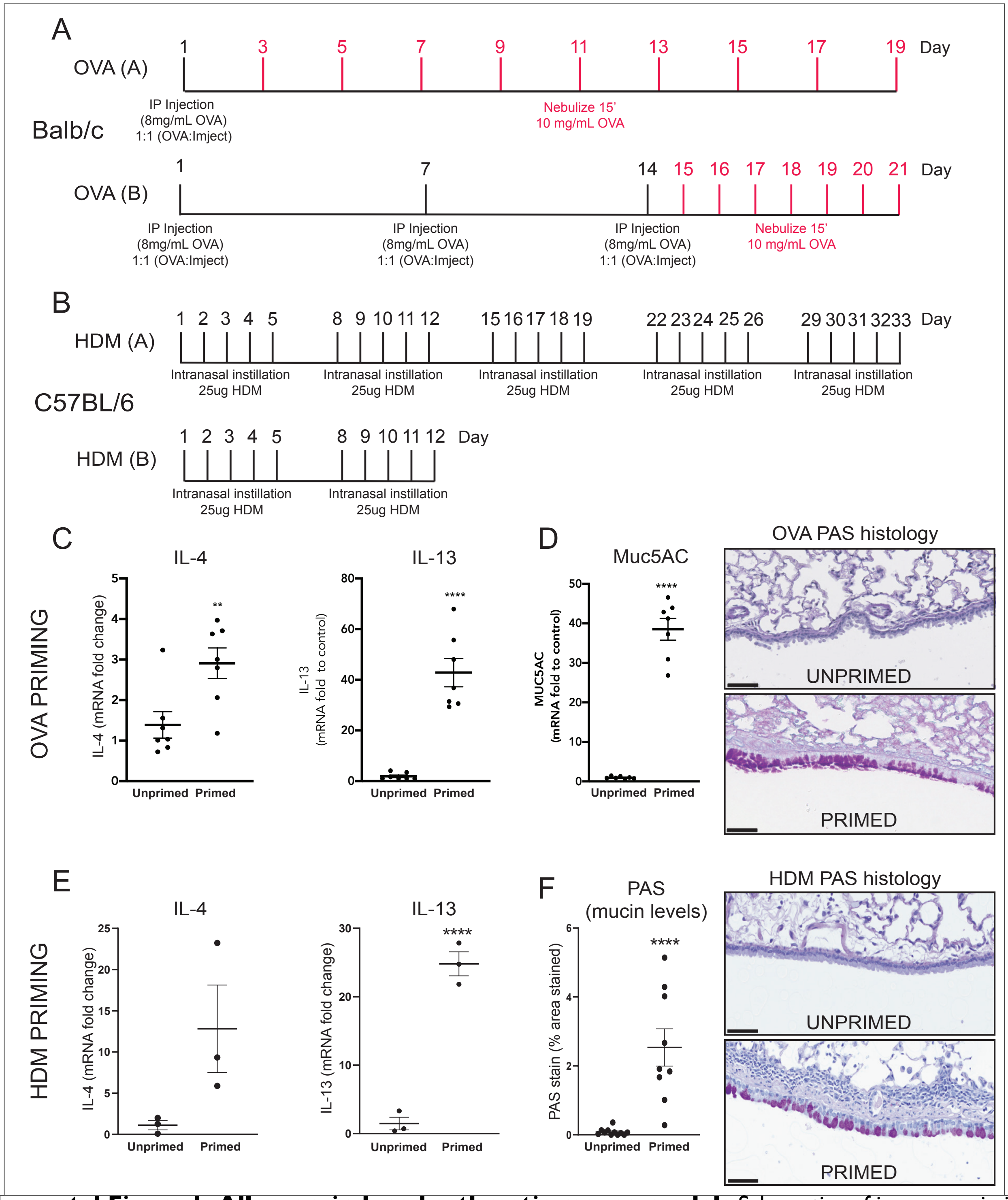


**Supplemental Figure 1. Allergen-induced asthmatic mouse models** Schematics of immune priming protocols for Balb/c OVA:Imject (A) and C57BL/6J HDM (B) asthmatic mouse models. All priming protocols produced a characteristic Th2 inflammatory response where IL-4- and IL-13 mRNA was analyzed by qRT-PCR from OVA B (C) and HDM A (E) primed mice, with at least 3 mice per group and P**<0.005, P****<0.0001. Immune priming resulted in excessive mucus production with increased mRNA levels of the pathological mucin Muc5AC (D) and increase PAS staining (F) from histological sections in OVA A (D) and HDM A (F) primed mice.

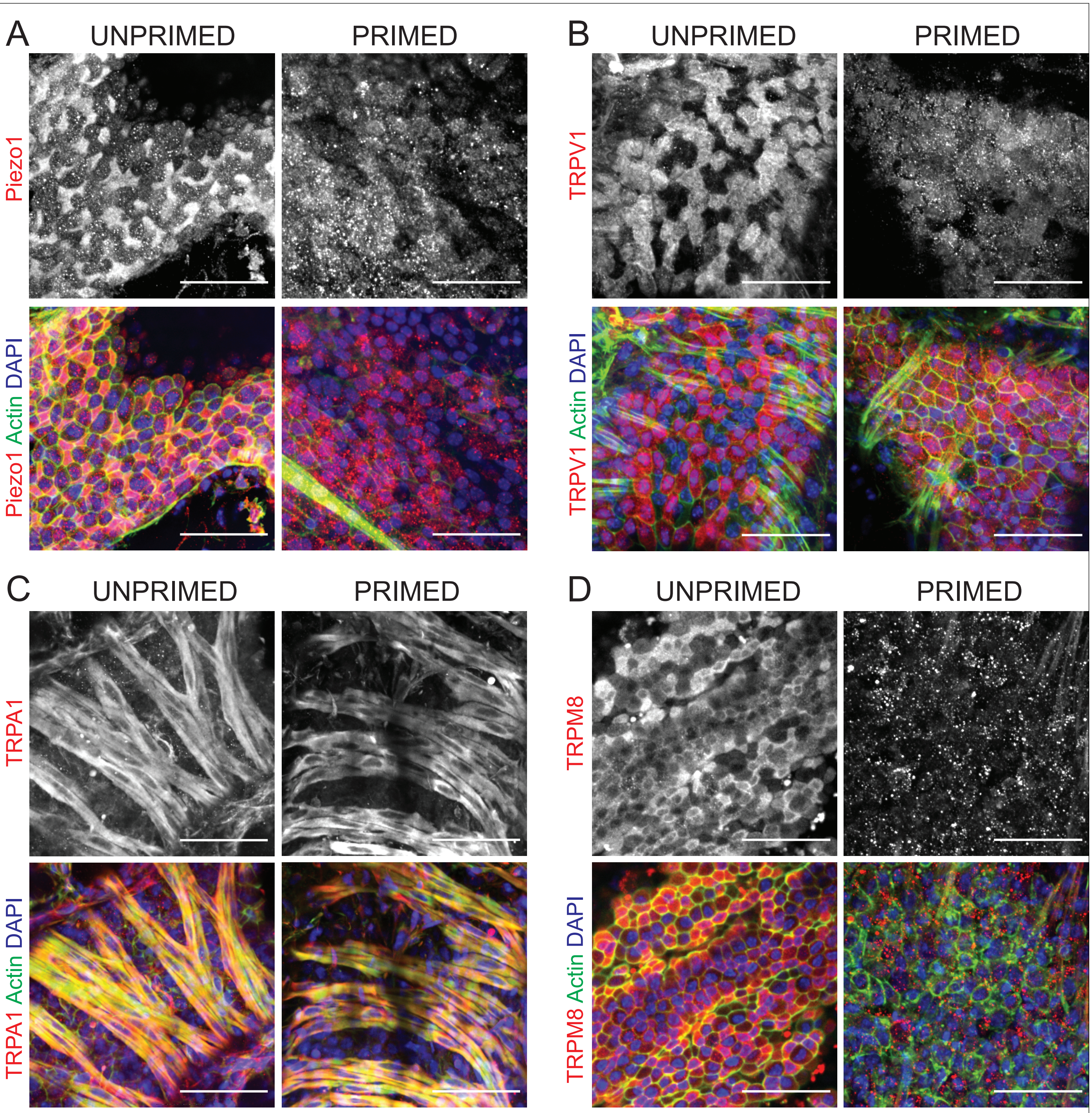


**Supplemental Figure 2. Stretch-activated and Transient Receptor Potential channels in airway epithelia** Representative confocal projections of airways from unprimed (Ctrl) and HDM-primed mice of channels that can be inhibited by gadolinium treatment: Piezo1 (A), TRPV1 (B), TRPA1 (C), and TRPM8 (D) (scale bars=50μm) from >3 mice per group.

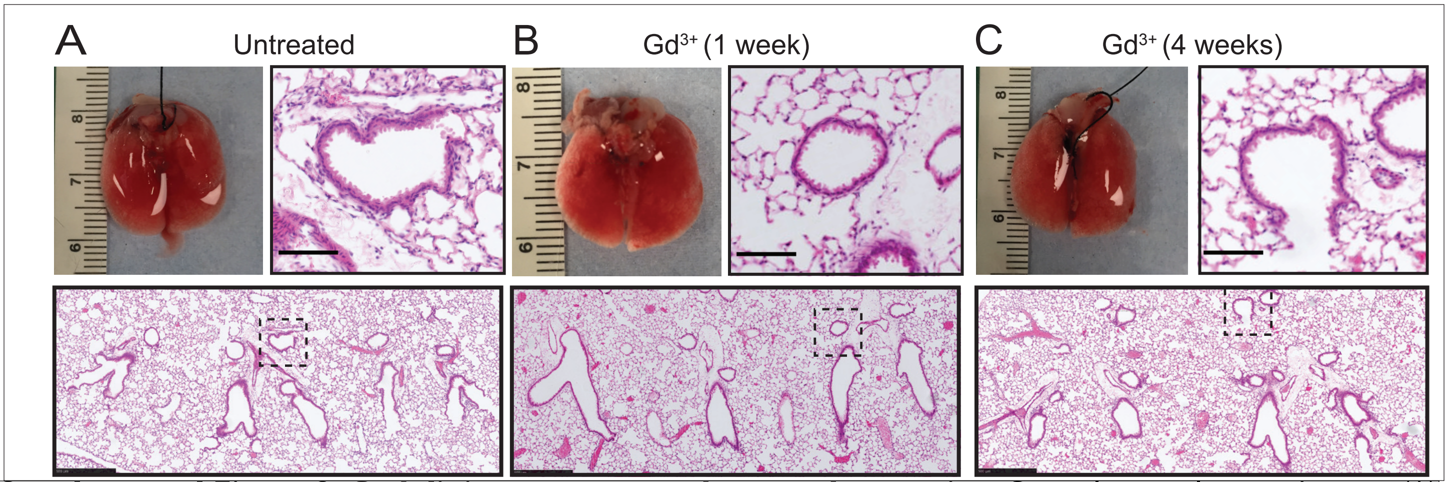


**Supplemental Figure 3. Gadolinium treatment does not harm mice** Control mouse lungs and tissues (A) or those from mice with intranasal instillation with 10μM $Gd^{3+}$ for 5 consecutive days (B) or once per week for 4 weeks (C) analyzing gross lung morphology and lung pathology by H&E staining. No discernable differences were noted between the groups (inset scale bar=100μm, whole image scale bar=500μm) from 3 lung slices per mouse from 4 mice per group.

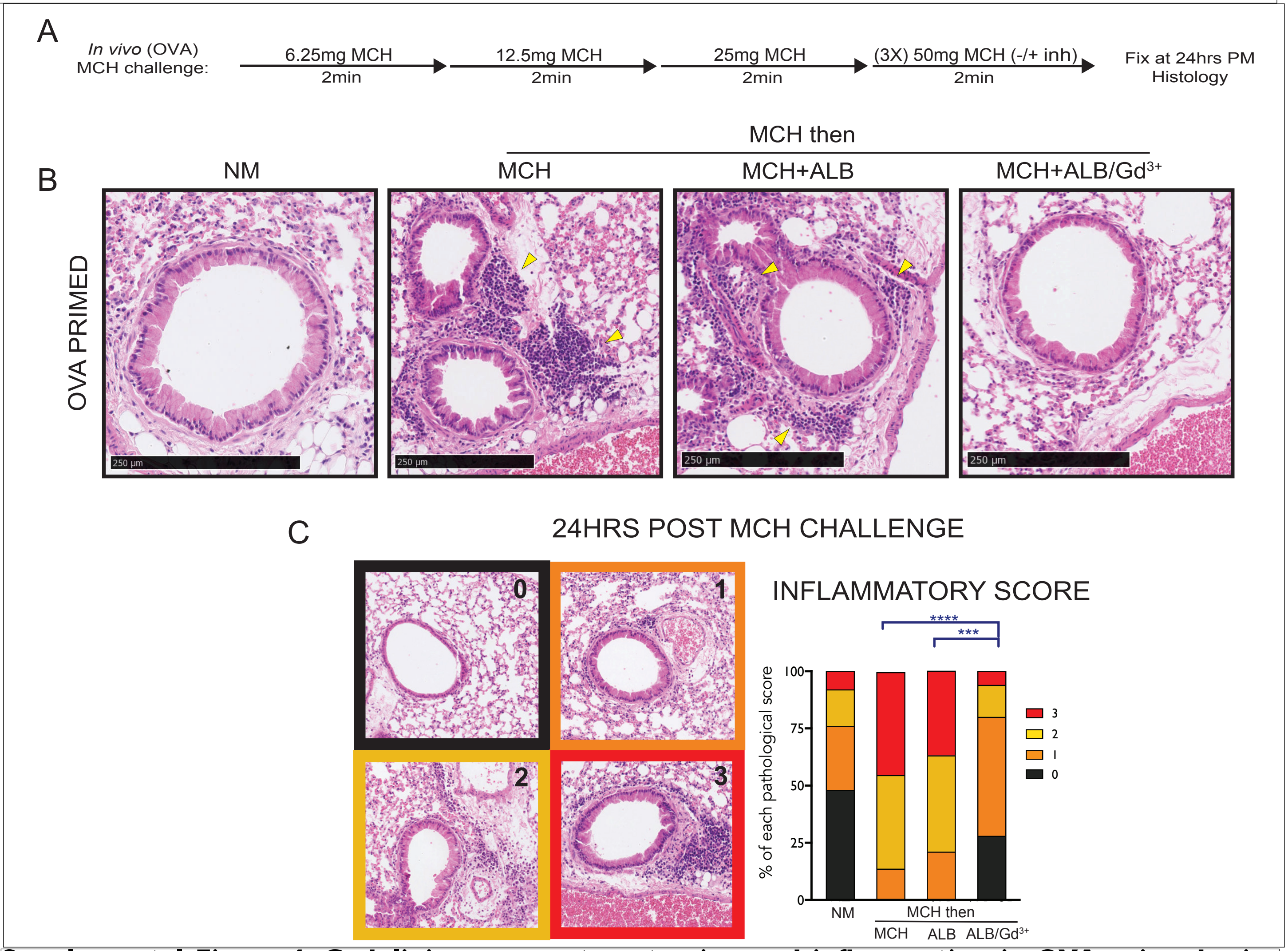


**Supplemental Figure 4. Gadolinium prevents extrusion and inflammation in OVA-primed mice** (A) Schematic of in vivo MCH challenges ± ALB or ALB/$Gd^{3+}$ in OVA primed mice. (B) Representative H&E sections from lung fixed at 24hrs after live mice were treated with: no MCH or with MCH ramp for 15min, then MCH, MCH+ALB, or MCH+ALB/$Gd^{3+}$, with yellow arrowheads indicating regions of high inflammatory cell infiltrates (scale bar=250μm), quantified in (C) from 5 mice from each treatment, with P***<0.0005 and P****<0.0001 from Chi-square tests.

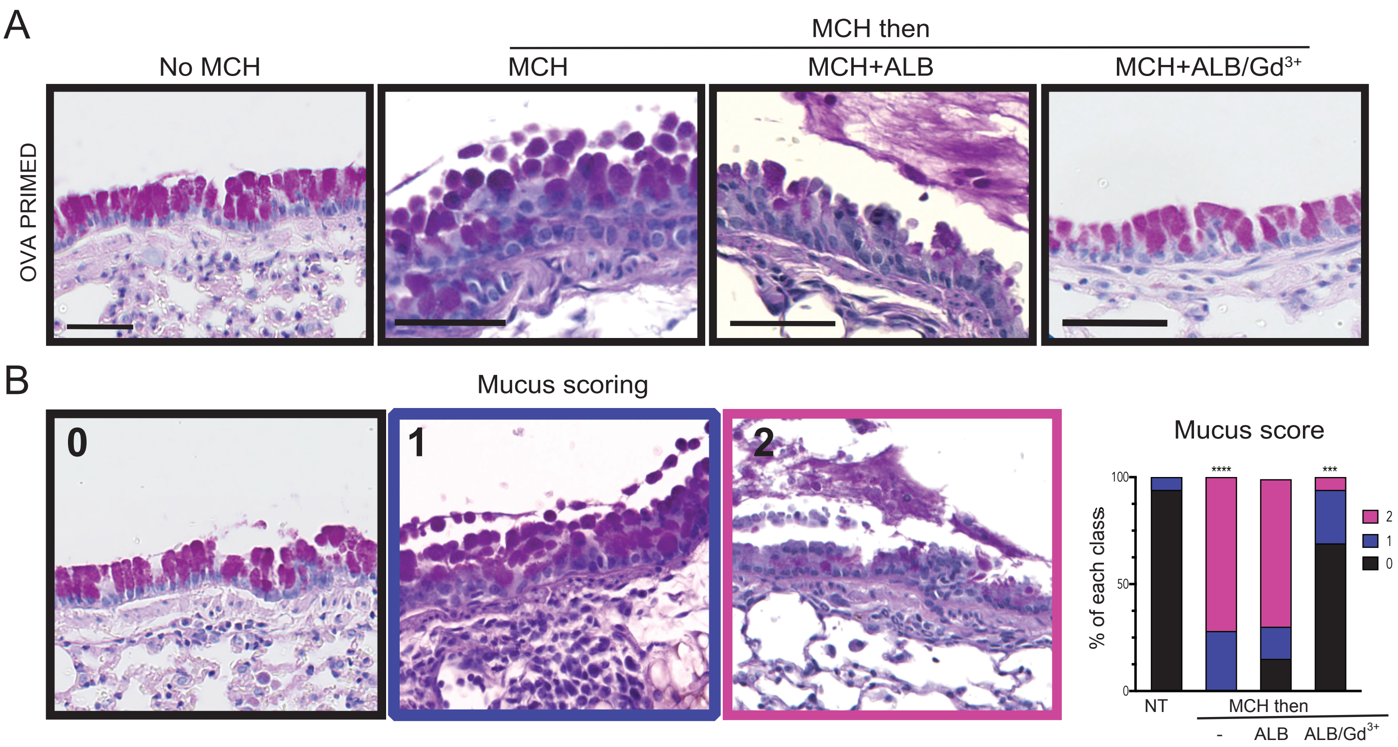


**Supplemental Figure 5. Gadolinium prevents MCH-induced mucus secretion in live OVA-primed mice** (A) Representative PAS stained sections from OVA-primed mice that were not treated with MCH or with MCH for 15mins before treating with MCH, MCH+ALB, or MCH+ALB/$Gd^{3+}$, quantified (as in Figure 4D) with a color/number score in (B). From 5 mice per treatment group with P***<0.0005 and P****<0.0001 from Chi-square tests

### UPPLEMENTARY FIGURES

**Supplemental Figure 1. Allergen-induced asthmatic mouse models** Schematics of immune priming protocols for Balb/c OVA:Imject (A) and C57BL/6J HDM (B) asthmatic mouse models. All priming protocols produced a characteristic Th2 inflammatory response where IL-4- and IL-13 mRNA was analyzed by qRT-PCR from OVA B (C) and HDM A (E) primed mice, with at least 3 mice per group and P**<0.005, P****<0.0001. Immune priming resulted in excessive mucus production with increased mRNA levels of the pathological mucin Muc5AC (D) and increase PAS staining (F) from histological sections in OVA A (D) and HDM A (F) primed mice.

**Supplemental Figure 2. Stretch-activated and Transient Receptor Potential channels in airway epithelia** Representative confocal projections of airways from unprimed (Ctrl) and HDM-primed mice of channels that can be inhibited by gadolinium treatment: Piezo1 (A), TRPV1 (B), TRPA1 (C), and TRPM8 (D) (scale bars=50μm) from >3 mice per group.

**Supplemental Figure 3. Gadolinium treatment does not harm mice**

Control mouse lungs and tissues (A) or those from mice with intranasal instillation with 10μM $Gd^{3+}$ for 5 consecutive days (B) or once per week for 4 weeks (C) analyzing gross lung morphology and lung pathology by H&E

staining. No discernable differences were noted between the groups (inset scale bar=100μm, whole image scale bar=500μm) from 3 lung slices per mouse from 4 mice per group.

**Supplemental Figure 4. Gadolinium prevents extrusion and inflammation in OVA-primed mice**
(A) Schematic of in vivo MCH challenges ± ALB or ALB/$Gd^{3+}$ in OVA primed mice. (B) Representative H&E sections from lung fixed at 24hrs after live mice were treated with: no MCH or with MCH ramp for 15min, then MCH, MCH+ALB, or MCH+ALB/$Gd^{3+}$, with yellow arrowheads indicating regions of high inflammatory cell infiltrates (scale bar=250μm), quantified in (C) from 5 mice from each treatment, with P***<0.0005 and P****<0.0001 from Chi-square tests.

**Supplemental Figure 5. Gadolinium prevents MCH-induced mucus secretion in live OVA-primed mice** (A) Representative PAS stained sections from OVA-primed mice that were not treated with MCH or with MCH for 15mins before treating with MCH, MCH+ALB, or MCH+ALB/$Gd^{3+}$, quantified (as in Figure 4D) with a color/number score in (B). From 5 mice per treatment group with P***<0.0005 and P****<0.0001 from Chi-square tests

**SUPPLEMENTARY MOVIES**

**Supplemental movie 1. Unprimed bronchioles mildly respond to MCH treatment.** *Ex vivo* lung slice from a healthy control mouse treated with 0.5g/mL MCH and imaged live by brightfield for 15 minutes (Fig. 1A).

**Supplemental movie 2. HDM priming airways results in pathological crowding and extrusion.** Lung slice from HDM B primed mouse was imaged live for 15mins of a 0.5g/mL MCH challenge (Fig. 1A).

**Supplemental movie 3. OVA priming airways results in pathological crowding and extrusion.** Lung slice from OVA A primed mouse was treated with 0.5g/mL MCH and imaged live for 30mins.

**Supplemental movie 4. Albuterol does not reverse epithelial cell extrusion during an attack.** A medium size airway from a five-week HDM-primed mouse treated with MCH for 15’, then MCH with albuterol (ALB) for an additional 15’ (Fig. 1F).

**Supplemental movie 5. Gadolinium protects airway epithelium even after an attack begins.** Airways from a five-week HDM-primed mouse treated with MCH for 15’ and then with MCH+ALB+$Gd^{3+}$ for another 15’. Red arrowheads pointing to areas of epithelial detachment, then $Gd^{3+}$-induced reattachment by the end of the treatment (Fig. 2E).

**Supplemental movie 6. Gadolinium protects airway epithelium even after an attack begins.** The uncropped Supplemental Movie 5 & Fig. 2E.

**Supplemental movie 7. Gadolinium does not appear to negatively impact mouse behavior.** Mice acutely treated with gadolinium on day 1 demonstrate normal grooming and social behavior.

**Supplemental movie 8. Gadolinium does not appear to negatively impact mouse behavior.** Mice acutely treated with gadolinium on day 7 demonstrate normal grooming and social behavior.

**Supplemental movie 9. Mechanically-induced mucus secretion in *ex vivo slices* by live WGA-350 imaging.** Inset of a bronchiole from a three-week HDM-primed lung pre-treated with 10µg/mL WGA before imaging for 15’ with 0.5g/mL MCH, showing mucus (as indicated by blue WGA) secreted from the epithelium in response to bronchoconstriction.

**Supplemental movie 10. Mechanically-induced mucus secretion in *ex vivo slices* by live WGA-350 imaging.** Uncropped Supplemental Movie 9 where a bronchiole from a three-week HDM-primed lung was pre-treated with 10µg/mL WGA before imaging for 15’ with 0.5g/mL MCH, demonstrating mucus (blue WGA) secreted from the epithelium in response to bronchoconstriction.

**Supplemental movie 11. Gadolinium pretreatment prevents mechanically induced mucus secretion.** *Ex vivo* lung slices from three-week HDM-primed mice were pretreated with WGA and $Gd^{3+}$ (10mins) then given 0.5g/mL MCH (t=00:00) and imaged live for 15’, demonstrating that most mucus is retained in the epithelium during bronchoconstriction.
